# haCCA: Multi-module Integrating of spatial transcriptomes and metabolomes

**DOI:** 10.1101/2024.08.20.608773

**Authors:** Xiao-Tian Shen, Jing Xu, Chen Zhang, Xiao-Yun Zhang, Zhou-Qing Chen, Hu-Liang Jiang, Lu-Yu Yang

## Abstract

Spatial techniques such as spatial transcriptomes and MALDI-MSI, offering insights into both transcripts and metabolite of tissue sections. However, integrating them with high accuracy is challenge due to no shared spots or features. We present haCCA, a workflow designed to integrate spatial transcriptomes and metabolomes data using high-correlated feature pairs and modified spatial morphological alignment. This approach ensures high-resolution and accurate spot-to-spot data integration across neighbor tissue section. We applied haCCA to both publicly available 10X Visium and MALDI-MSI datasets from mouse brain tissue and a custom spatial transcriptome and MALDI-MSI dataset from an intrahepatic cholangiocarcinoma (ICC) model, exploring the metabolic alteration of NETs(neutrophil extracellular traps) on ICC, and finding a potential mechanism that NETs upregulated Scd1 to activate fatty acid metabolism. Providing new insights into the dynamic crosstalk between genes and metabolites that regulates the tumor biological behavior and drives the response to treatment. We developed and published an easy-to-use Python package to facilitate its use.

## Introduction

Spatial multiomics techniques allows resolving the spatial molecular profiles at transcript, metabolite, protein or epi-genetic level[1]. Spatial transcriptomes enable the measurement of mRNA expression, while MALDI-MSI(Marix-Assisted Laser Desorption Ionization Mass Spectrometry Imaging) allows the collection of mass-to-charge (m/z) spectra at defined raster spots across the tissue section. Both methodologies are most frequently applied spatial techniques. These 2 techniques generate gene-expression or peak-intensity matrix of sampling spots, with coordinates indicating their location. Numbers of spots can vary from thousands to hundreds of thousands determined by resolution.

An integration of spatial transcriptomes and MALDI-MSI to generate multi-module spatial data provides more comprehensive understanding and novel insight to biological state of section spots. However, such integration may be challenge. Till now 3 categories of integration strategies has been brought out for multi module single cell data[2], using mutual data features, using mutual cells, or using shared latent space if both mutual cells or features are not accessible. The first 2 strategies are able to keeps the origin resolution, which is single cell or spot, while the last strategy will lose it. Usually, no mutual feature can be found between spatial transcriptomes and MALDI-MSI, neither mutual spots or cells, constrain the application of first 2 strategies. Luckily, spatial multi module techniques are often applied on neighbor sections, even the same section[3], ensuring the morphological similarity. Hence morphological alignment of spatial shape followed by nearest neighbor spot pair finding strategy has been applied on their integration most often[4-7].

Such integrate strategy brings certain flaws. First is that there is no benchmark data to access the performance of integration, second is that as no features are applied for assisting the integration in present procedure, the efficiency of integration might be undermined. Using mutual features for alignment has been proved useful in integrate single cell data from difference batches, where by using linear combination of mutual features to establish latent space, allowing cells with the same labels to appear together and be clustered into the same group [8-10]. Although no mutual feature existed among MALDI-MSI and spatial transcriptomes, a few studies revealed the high correlation between gene transcripts and metabolites[11, 12], giving us the inspiration to explore whether high correlated gene-metabolite pairs can be utilized in integrating.

In this study, we established haCCA, a workflow utilizing **h**igh Correlated fe**a**ture pairs **c**ombined with a modified spatial morphologi**c**al **a**lignment to ensure high resolution and accuracy of spot-to-spot data integration of spatial transcriptomes and metabolomes. We generated a series of benchmarks using spatial transcriptomes and paired pseudo spatial metabolomes, proving haCCA outperformed morphological-aligned method(for example, STUtility). We tested the performance of haCCA in assisting multi-module spatial data analysis using public available 10X Visium and MALDI-MSI data cohort of mouse brain and a home-brew spatial transcriptomes(on BMKMANU S1000 platform) and MALDI-MSI data cohort of ICC(intrahepatic cholangiocarcinoma) model. Using haCCA, we performed an in situ and in vivo analysis and found the metabolic alteration effect of NETs(neutrophil extracellular traps) on ICC. NETs upregulated Scd1 to activate multiple metabolism, especially fatty acid elongation. An easy-to-use python package has been developed and published for use.

To summarize, we presented a workflow named haCCA that enables the simultaneous spatial profiling of metabolites and transcriptome across neighbor tissue section. Applying haCCA to tumor samples could increase understanding of both metabolic and transcript heterogeneous, providing new insights into the dynamic crosstalk between genes and metabolites that regulates the tumor biological behavior and drives the response to treatment.

## Method

### Data source

For public data, 10X Visium and MSI-MALDI data from Parkinson’s disease mouse brain was download from Macro et al’s manuscript[3]. 6 10X Visium,3 MSI-MALDI data of 6 sections from 3 mouses were included(table S1).

For home-brew data, We generated Akt/Yap induced ICC(intrahepatic cholangiocarcinoma) mouse model in WT or Padi4-/-mouse. BMKMANU S1000 Spatial transcriptomics and MALDI-MSI data of tumor was generated following standard procedure (see below).

### Mouse model

6-8 weeks old of male C57BL/6 mouse were purchased from https://www.gempharmatech.com/ and kept in a SPF institute at 25°C temperature, 50-60% moisture. Pre-clinical ICC model was generated by HDI (hydrodynamic injection) of 20ug pT3-myrAkt-p2a-Yap(S127A) and 5ug S100B plasmid in 6w old wt or Padi4-/-C57 mouse. Padi4 -/-mouse is a kind gift from Prof Chen-De Yang. After 4 weeks, tumors were collected for further experiment. (Figure S1)

### BMKMANU S1000 Spatial transcriptomics

#### Frozen embedded tissue

After tissue samples was obtained, tissue surfaces were quickly rinsed with a pre-cooled solution of 1X PBS (RNase free) or normal saline to remove residual blood, and sterile gauze was used to blot the surface fluid. The tissue size was required to be suitable, and the tissue should be cut into small pieces(6.8mm2) suitable for subsequent experiments. Small fragments of each tissue were snap-frozen in isopentane pre-chilled with liquid nitrogen and optimum cutting temperature (OCT) compound (SAKURA,Cat#: 4583) and stored at −80°C until use.

#### Slide preparation

Spatial Transcriptomics slides were printed with 1-8 identical 6.8×6.8 mm capture areas, each with 2,000,000 spots contain barcoded primers (BMKMANU S1000). The primers are attached to the slide by the 5’ end and contain a cleavage site, a T7 promoter region, a partial read1 Illumina handle, a spot-unique spatial barcode, a unique molecular identifier (UMI), and Poly(dT)VN. The spots have a diameter of 2.5μm and are arranged in a centered regular hexagonal grid so that each spot has six surrounding spots with a center-to-center distance of 4.8μm. The frozen tissue was cut in a pre-cooled cryostat at 10µm thickness and systematically placed on chilled BMKMANU S1000 Tissue Optimization Slides and BMKMANU S1000 Gene Expression Slides, and stored at −80°C until use.

#### Tissue optimization

The Spatial Transcriptomics (ST) protocol was optimized for tissue according to recommendations. In short, changes were made in the staining procedure by excluding isopropanol, decreasing the incubation time of hematoxylin and bluing buffer, as well as increasing eosin concentration. Moreover, the optimal incubation time for permeabilization was established, and the previously described one-step protocol for tissue removal was altered by using a higher proteinase K:PKD buffer ratio. Once optimal conditions had been established, three cryosections per patient were cut at 10 mm thickness onto spatial slides and processed immediately.

#### Fixation, staining and imaging

Sectioned slides were incubated at 37°C for 1 min., fixed in 3.7%–3.8% formaldehyde (Sigma-Aldrich) in PBS (Medicago) for 30 min, and then washed in 1x PBS (Medicago). For staining, sections were incubated in Mayer’s hematoxylin (Dako, Agilent, Santa Clara, CA) for 4 min, bluing buffer (Dako) for 30 s, and Eosin (Sigma-Aldrich) diluted 1:5 in Tris-base (0.45M Tris, 0.5M acetic acid, pH 6.0) for 30 s. The slides were washed in RNase and DNase free water after each of the staining steps. After air-drying, the slides were mounted with 85% glycerol (Merck Millipore, Burlington, MA) and coverslips (Menzel-Glaser). Bright-field (BF) images were taken at 20x magnification using Metafer Slide Scanning platform (MetaSystems). Raw images were stitched with VSlide software (MetaSystems). The coverslip and glycerol were removed after imaging by immersing slides in RNase and DNase free water. The slides were inserted into slide cassettes to separate the tissue sections into individual reaction chambers (hereinafter wells). For pre-permeabilization, sections were incubated at 37°C for 20 min with 0.5 U/ml collagenase (ThermoFisher) and 0.2 mg/ml BSA (NEB, Ipswich, MA) in HBSS buffer (ThermoFisher). Wells were washed with 0.1× SSC(Sigma-Aldrich), after which permeabilization was conducted at 37°C for 7 min in 0.1% pepsin (Sigma-Aldrich) dissolved in 0.1M HCl (Sigma-Aldrich). After incubation, the pepsin solution was removed and wells washed with 0.1 × SSC.

#### Reverse transcription, spatial library preparation and sequencing

Reverse transcription (RT), second-strand cDNA synthesis, adaptor ligation and a second RT was generated and libraries were constructed according to the performer’s protocol. Sequencing handles and indexes were added in an indexing PCR and the finished libraries were purified and quantified. Sequencing was performed on the Illumina NovaSeq 6000 with a sequencing depth of at least 50,000 reads per sopt(100μm) and 150bp (PE150) paired-end reads (performed by Biomarker Technologies Corporation, Beijing, China).

#### Spot visualization and image alignment

Primer spots were stained by hybridization of fluorescently labeled probes and imaged on the Metafer Slide Scanning platform. The resulting spot image was loaded into the BSTMatrix and BSTViewer along with the previously obtained BF tissue image of the same area. The two images were aligned and the built-in tissue recognition tool was used to extract spots covered by tissue. 7

#### BSTMatrix analysis

Finished libraries were diluted to 4nM and sequenced on the Illumina Nova 6000 using paired-end sequencing. We completed the upstream analysis through BSTMatrix (v1.0). The mapping was performed to the reference GRCh38_release95 mouse genome.

#### Downstream analysis

Briefly, data analysis was carried out through steps of normalization, clustering and screening marker genes using Python package pyscenic.

### MALDI-MSI

#### Matrix coating

Desiccated tissue sections mounted on ITO glass slides were sprayed using an HTX TM sprayer (Bruker) with 10 mg/mL 9AA (9-aminoacridine), dissolved in ethanol-water (7:3, v/v). The sprayer temperature was set to 90°C, with a flow rate of 0.12 mL/min, pressure of 10 psi. Four passes of the matrix were applied to slides with 10 s of drying time between each pass.

#### Mass spectrometry imaging

MALDI timsTOF MSI experiments were performed on a prototype Bruker timsTOF flex MS system (Bruker) equipped with a 10 kHz smart beam 3D laser. Laser power was set to 70% and then fixed throughout the whole experiment. The mass spectra were acquired in negative mode. The mass spectra data were acquired over a mass range from *m/z* 50-1200 Da. The imaging spatial resolution was set to 30 μm for the tissue, and each spectrum consisted of 400 laser shots. MALDI mass spectra were normalized with the Root Mean Square, and the signal intensity in each image was shown as the normalized This experiment was performed at biotree company(www.biotree.cn).

#### Pseudo-MALDI-MSI data generating

We employed an iteration and sampling strategy on spatial transcription data to generate paired pseudo-MALDI-MSI data. These pairs served as benchmarks for evaluating the performance of our alignment strategy.

For the spatial transcription data, we have 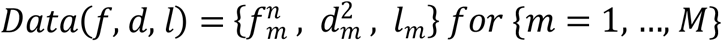 *for* {*m* = 1, …, *M*}, where for each *data*_*m*_(*f, d, l*) f denotes the n-dimension feature-spot vector *f*^1^, *f*^2^, …, *f*^*n*^}, d represents the coordinate matrix (*d*_*x*_, *d*_*y*_), *l* represents the cluster label.

Given *Data*(*f,d,l*) we use the following steps to mimic the morphological disagreement between neighboring sections and generate a pair of 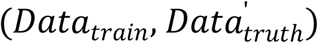 from *Data*(*f,d,l*)to evaluate *haCC****A*** workflow.

**Step 1:** Split feature set *f* by randomly sample *j where j* ∈ (0, *n*)features from *f* and assign them to 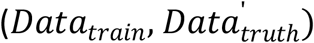

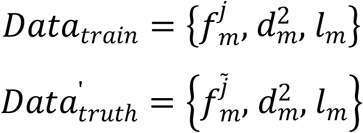

**Step 2:** apply Gaussian translation to both *d*_*train*_ and *d*_*truth*_ to mimick the morphological disagreement between neighborsections.

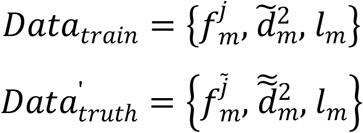

#### haCCA workflow

Given two spatial data ***Data***_***A***_ and ***Data***_***B***_ where

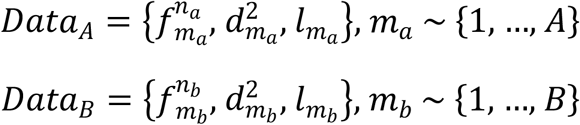

***haCCA***_***A***→***B***_ aligns the points in ***Data***_***A***_ to ***Data***_***B***_ by finding a point-to-point map from *A* to *B* which minimizing the following loss object

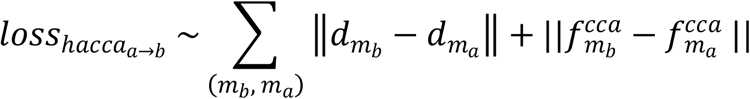

Where {(*m*_*b*_,*m*_*a*_),*m*_*b*_ ∈*Data*_*b*_, *m*_*a*_ ∈ *Data*_*a*_ } is the aligned points pairs. Once {(*m*_*b*_,*m*_*a*_)} is determined, the result of *haCCA*_***A***→***B***_ and aggregate result for can be represented as

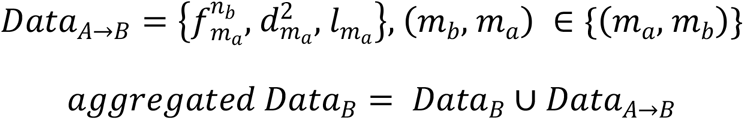

To minimize the 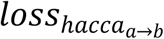 we performed a heuristic two-step method: In the first step, we apply the gross alignment plus further alignment to *Data*_***B***_ which minimize the first item of 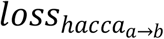. In the second step, we make use the alignment result from first step and identify a set of anchor points, which is used to determine the high correlation feature pairs and minimize the second item of the loss function. We will introduce each step in detail in the below sections.

**Step 1: minimize the first item by aligning A to B using location information {*d*}**

In this step, we apply two alignment strategies here: gross alignment and further alignment.

In gross alignment, manual alignment or classical cloud mesh alignment algorithm like ICP or FGW is used to determine an affine translation matrix *M* to minimize the L2 distance between points in *Data*_***A***_ and *Data*_***B***_

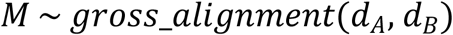

*d*_*B*_= *M* ∗*d*_*B*_ manual alignment is processed using the following steps:

1) manually select *n* points from *m*_*a*_ and *m*_*b*_, denoted as 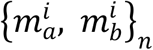.
2) Calculate the affine translation matrix *M* using the location of 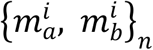.

In further alignment, we find a rotation and moving matrix *M’* to minimize the distance between outlier spots in {*d*_*B*_} and their nearest neighbors in the outlier spots in {*d*_*B*_} is defined as the spots where it does not have neighbor within a given distance in {*d*_*A*_}. After the outlier spots in {*d*_*B*_} is found, we calculate the affine translation matrix *M’* using outlier spots in {*d*_*B*_} and its nearest neighbor in {*d*_*A*_}.

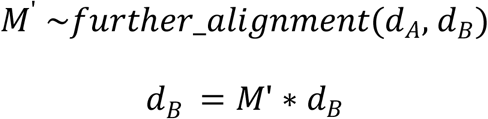

**Step 2: minimize the second item by reducing the distance of high-correlation feature pairs identified by anchor spot pairs** 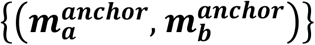, *m*_*a*_ ∈ *M*_*A*_, *m*_*b*_ ∈ *M*_*B*_

The anchor spot pair are paired points from *(M*_*A*_, *M*_*B*_) where each pair *(m*_*a*_, *m*_*b*_) satisfies with the following conditions: 1. The distance between *m*_*a*_ and *m*_*b*_ are smaller than a given threshold, and they are the closest point to each other. 2. The neighborhood of both *m*_*a*_ and *m*_*b*_ is of low variety. We use Simpson index to evaluate the variety.

The anchor spot pairs are identified using the following steps: 1. For each points {m_a_} ∈ *M*_*A*_, found a set of points {*m*_*b*_},*m*_*b*_ ∈ *M*_*B*_ where each *m*_*b*_ is within a given distance. 2. For each item in {*m*_*a*_: {*m*_*b*_} }, *m*_*a*_ ∈ *M*_*A*_, *m*_*b*_ ∈ *M*_*B*,_ calculate the Simpson index for each item and remove the items where its Simpson index is higher than a given threshold. 3. For the remaining items in {*m*_*a*_: {*m*_*b*_} }, *m*_*a*_ ∈ *M*_*A*_,*m*_*b*_ ∈ *M*_*B*,_the final anchor spot pairs 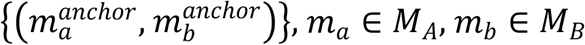 are identified as the closest point pair in each item.

After finding anchor spot pairs 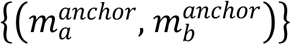, *m*_*a*_ ∈ *M*_*A*_, *m*_*b*_ ∈ *M*_*B*,_ the high correlated feature pairs 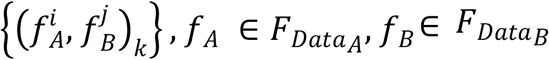 where *i,j* identifies the *i*^*th*^and *j*^*th*^feature in each dataset are further identified by the top *k* score of Pearson correlation matrix on aggregated feature sets 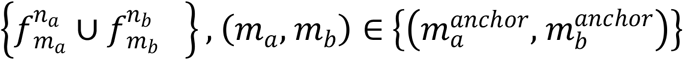

After the high correlated feature pairs 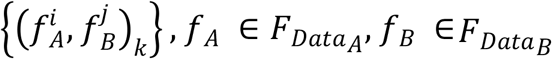 identified, we calculate the CCA feature for both {*m*_*a*_} and {*m*_*b*_} using canonical variables found by Canonical Correlation Analysis (CCA). The idea behind is to further maximize the correlation and normalize the value between the high correlated feature pairs. The new features are noted as *f*^*cca*^. *f*^*cca*^ will be added to *Data*_*A*_ and *Data*_*B*_ as an individual feature after they are calculated for all the points {*m*_*a*_} in *Data*_*A*_ and {*m*_*b*_} in *Data*_*B*_.

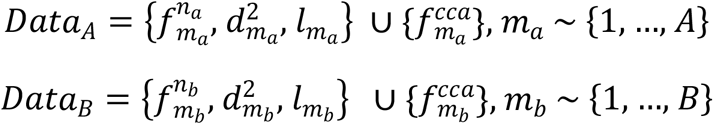

Finally, we minimize the second item on both {*l*} and {*f*^*cca*^} using classic cloud mesh alignment algorithm like ICP or FGW.

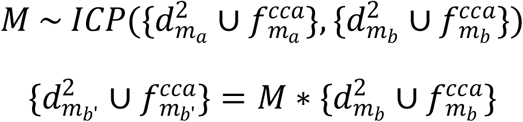

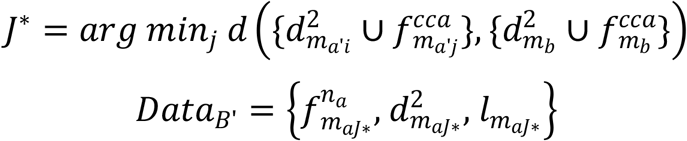

*FGW* algorithm is described below:

Finding a transport plan π, where

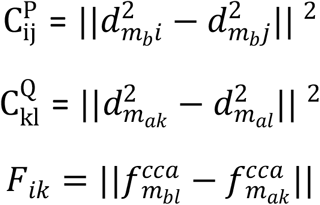

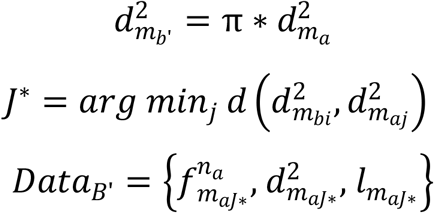

B’ is the haCCA translated B.

### Evaluation

#### Label Transfer Accuracy

Label transfer accuracy measures how well labels from one dataset can be transferred to and correctly applied in another dataset.

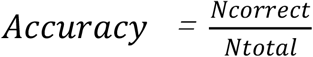

Where N_correct_ is the number of correctly transferred labels in the target dataset, N_total_ is the total number of samples in the target dataset.

#### Label Transfer ARI

Label Transfer ARI (Adjusted Rand Index) is a measure of the similarity between two data clusterings or label assignments. It adjusts the Rand Index (RI) to account for chance grouping.

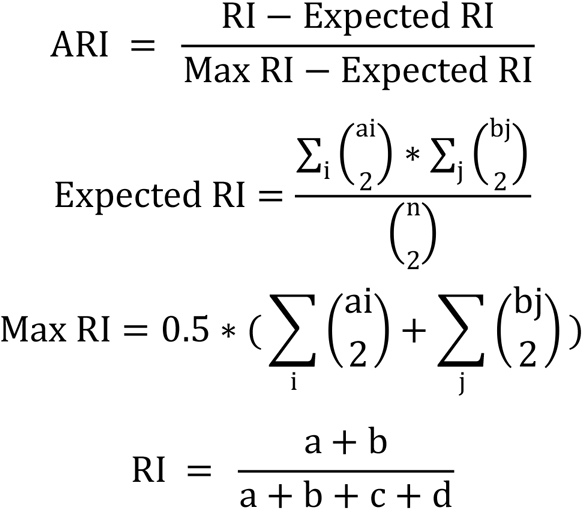

n: Total number of elements in the dataset.

a: Number of pairs of elements that are in the same cluster in both clusterings.

b: Number of pairs of elements that are in different clusters in both clusterings.

c: Number of pairs of elements that are in the same cluster in the first clustering but in different clusters in the second clustering.

d: Number of pairs of elements that are in different clusters in the first clustering but in the same cluster in the second clustering.

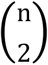: Total number of possible pairs of elements, calculated as n(n-1)/2.

#### Loss

For Data(X(m,n), D(m,2), Label(m,1)) and groundtruth Data’(X’(m,n), D’(m,2), Label’(m,1)), L2 difference between X and *X’ was* defined as Loss.

### Data analysis

Statistical analyses were performed using the R 4.0.4 software. Paired Student t test were used for comparisons between groups. For all tests, significance was determined with a 95% confidence interval (ns, P > 0.05; *, P < 0.05; **, P < 0.01; ***, P < 0.001; ****, P < 0.0001).

## Result

### The workflow of high Correlated feature pairs combined with spatial morphological alignment(haCCA) for Multi-module Spatial assay Integrating

haCCA integrates multi-module spatial assay by allocating spot data from one assay to the paired spot in another assay. After generating spatial transcriptome and MALDI-MSI data among neighbor section, haCCA integrates 2 assay through 6 steps: **1: data preparing**. Following the standard processing with Scanpy, the spatial data along with the peak/intensity or feature matrix are extracted, normalized, and scaled. **2: gross alignment:** Three feature points in both datasets, sharing high similarity in image characteristics, are manually selected to generate an affine matrix. This matrix enables the rotation, scaling, and translation of the MALDI-MSI in transcript spatial data to align with the spatial transcriptome data. **3: further alignment:** After gross alignment, a small subset of spots in spatial transcriptome I data are not overlay with MALDI-MSI. Moreover, the spot-spot distance between MALDI-MSI and spatial transcriptome is not minimized. We extract outlier spot in spatial transcriptome data, which is identified as having no neighbor spot in MALDI-MSI within certain distance.(dist_min). The nearest spot in MALDI-MSI for each outlier spot in spatial transcriptome data is identified as the potential neighbor spot. A translation matrix was found by minimizing the summary of distance between outliers and their potential neighbor spot. After this step, we found a satisfied alignment of the morphological shape between MALDI-MSI spatial data and spatial transcriptome data. **4: identification of anchor spot pairs and high correlated feature pairs**. After further alignment, we assume the morphological shape of MALDI-MSI and spatial transcriptome data are aligned. Anchor spot pairs between MALDI-MSI and spatial transcriptome data were admitted if they meet 2 criteria: 1. very close distance(default 0.5) 2. neighbor spots (distance less than certain value, default 2) are with low diversity (Simpson index of cluster <0.2). Anchor spots pairs are considered as spot appear at the same coordinates and their gene feature data from spatial transcriptome and metabolic data from MALDI-MSI are applied for pearson correlation test, in order to figure out high correlated feature pairs. **5: combined feature and morphological alignment**, paired features are undergoing CCA transfer to generate a high-correlated variate as a third dimension. Alignment using morphological data and high-correlated variate between MALDI-MSI and spatial transcriptome is carried out by ICP(Iterative Closest Point) algorithm. **6: integrating:** after ICP algorithm, MALDI-MSI and spatial transcriptome is projected to a 3-dimention, spatial-variable space. For each spot in spatial transcriptome data, nearest MALDI-MSI spot data is integrated to and generate multi-module data.(Figure 1)

**Figure 1.**
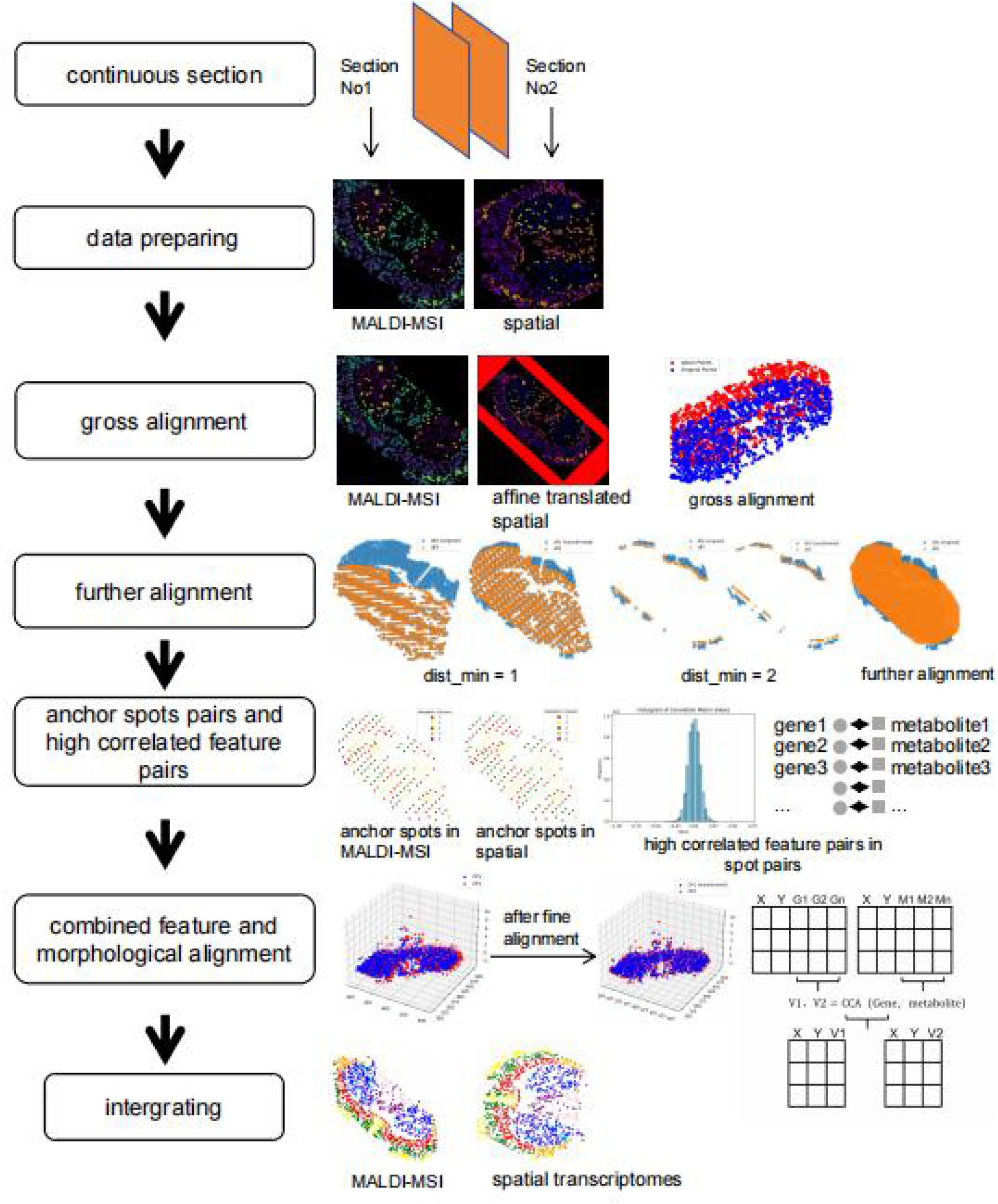
an overview illustrator of high Correlated feature pairs combined with spatial morphological alignment(haCCA). The procedure consist of 6 main steps: **Data preparing:** Peak-intensity in MALDI-MSI and normalized gene count in spatial transcriptome were processed and clustering with scanpy. **Gross alignment:** Feature points were selected based on mutual spatial region with high similarity between MALDI-MSI and spatial transcriptome data. Affine translation matrix was applied on Spatial matrix of spatial transcriptome data to achieve maximum similarity, shown in gross alignment. Red refers to MALDI-MSI and blue refers to affine translated spatial transcriptome data. **Further alignment:** after gross alignment, spots in spatial transcriptome within certain distance(dist_min) to MALDI-MSI were one-to-one paired aligned. If no paired spot was found, then such spots in spatial transcriptome was defined as outliers. Distance of outliers to nearest MALDI-MSI spots was calculated and a transfer matrix was found to minimize the sum. Then the transfer matrix was applied to the affine transferred spatial transcriptome. As shown, after further alignment a maxim alignment of morphological shape between MALDI-MSI and spatial transcriptome data was achieved. **Anchor spot pairs and high correlated feature pairs identification:** After further alignment, anchor spots, defined as spots pairs with the same location(distance < 0.1) were separately identified in spatial data of MALDI-MSI and spatial transcriptome data. Then high correlated features between anchor spots pairs was found by pearson correlation test. **Combine feature and morphological alignment:** Paired features are undergoing CCA transfer to generate a high-correlated variate as a third dimension. Alignment using morphological data and high-correlated variate between MALDI-MSI and spatial transcriptome is carried out by ICP(lterative Closest Point) algorithm, **integrating:** after ICP algorithm, MALDI-MSI and spatial transcriptome is projected to a 3-dimention, spatial-variable space. For each spot in spatial transcriptome data, nearest MALDI-MSI spot data is integrated to and generate multi-module data.

### haCCA perform effective integration on pseudo muti-module spatial data

To quantitatively evaluate the performance on integrating spatial transcriptome data and MALDI-MSI of haCCA, we generated pseudo MALDI-MSI data using transcript data. The pseudo MALDI-MSI has a few characters: 1. Small iteration was added to the morphological coordinates of pseudo MALDI-MSI data to ensure few morphological similarity with parent spatial transcriptome data. 2. no mutual features with parent spatial transcriptome data. We assessed label transfer accuracy (AC) of the cluster between ground truth and integrating outcomes to evaluate performance. In all, pseudo MALDI-MSI data was generated form 3 mouse brain, 1 rat brain and 1 human gastric cancer(n=15 each). First, we ask how each steps in haCCA contribute to integrating. We found that after gross and further alignment the AC increased significantly. Moreover, the combined feature and morphological alignment strategy also increased the AC from 6%-20%, depending on the AC in previous steps. This indicated the efficiency of combing strategy of haCCA (Fig2 A,B). Till now there is only very few public method to integrate multimodule spatial data. Most of them relied on morphological similarity. We found that compared with direct alignment, manual alignment(SUTility) and ICP, hCCA showed the best performance, with the highest CA and ARI(Fig 2C,D). These results indicating the efficiency of haCCA in integrating spatial transcriptome data and MALDI-MSI.

**Figure 2.**
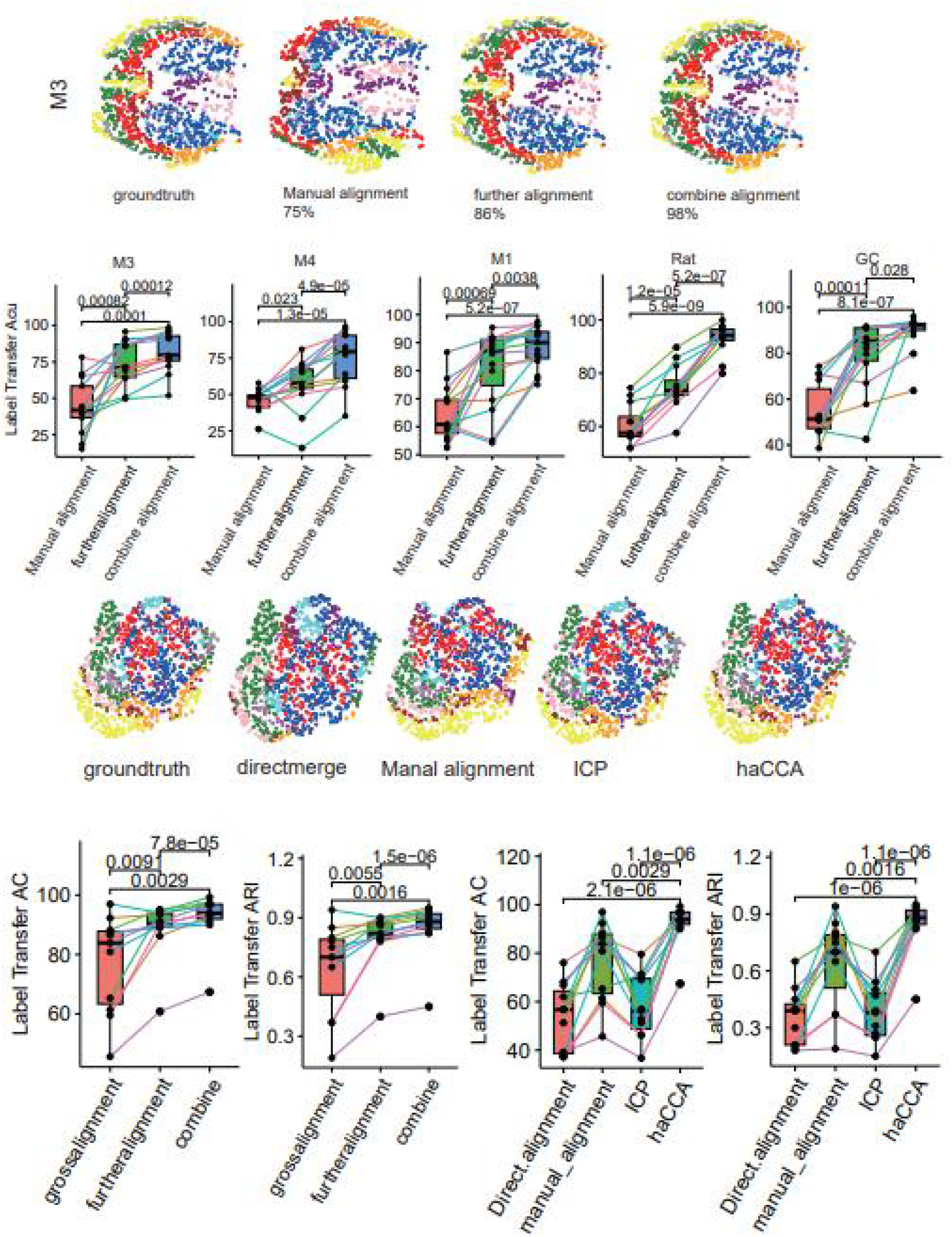
haCCA performance on pseudo muti-module spatial data A,B: Label Transfer Accuracy of 3 step of haCCA (gross alignment, Further alignment and combine morphological and feature pair alignment) on integrating spatial transcriptome and paired pseudo MALDI-MSI data. 5 batches were tested(n=15 each) and M3 batch was used as representation. C,D: comparation of label transfer accuracy and label transfer ARI between available integrating method. Such as directmerge, manual alignment(STUtility), ICP and haCCA. 5 batches were used for comparation and GC(gastric cancer) was used for representation.

### haCCA performs consistent integration of 10X Visium and MALDI-MSI spatial data

Next we asked how haCCA performed in real-world generated data. We applied haCCA on a dataset which is consisted of 10X Visium and MALDI-MSI from 3 mouse brain. For each mouse brain we included 3 spatial data from 2 section. To be noted, 1 run of 10X Visium and 1 run of MALDI-MSI data was performed on the same section, with a novel procedure. The third run of 10X Visium was performed on the neighbor section. STUtility was applied to integrated the 10X Visium and MALDI-MSI data carried on the same section by morphological alignment, which was used as the ground truth. We used haCCA to perform integration between 10X Visium data and MALDI-MSI data from the same section or located on the neighbor section. It turns out that haCCA is able to effectively integrated 10X Visium and MALDI-MSI spatial data from the same section, with the highly accuracy of 87.5%, 79.5% and 87.6%(Fig 3 A,C,E). as we applied STUtility outcomes as ground truth, and STUtility is of inferior performance in simulated data compared with haCCA, it is highly likely that haCCA outcomes is more accurate. In the mission of integrating 10X Visium and MALDI-MSI spatial data from neighbor section, where exists a larger difference between morphological shape. haCCA effectively transferred the morphological shape of 10X Visium data, making it more assemble to the MALDI-MSI data, and performed highly consistent integration, shown by the similar pattern of the metabolic cluster distribution between MALDI-MSI data and haCCA integrated data.(Figure 4 B,D,F) Using dopamine and 3-MT as example, it turns out haCCA is able to reconstruct the metabolite distribution in integrated data with the background of spatial transcriptome data (Fig 3G,H). It leverages the connivance to identify the genes associated with metabolite. Not surprisingly, genes associated with dopamine metabolism, such as Th, shown an overlapped distribution pattern with dopamine (Fig 3G).

**Figure 3.**
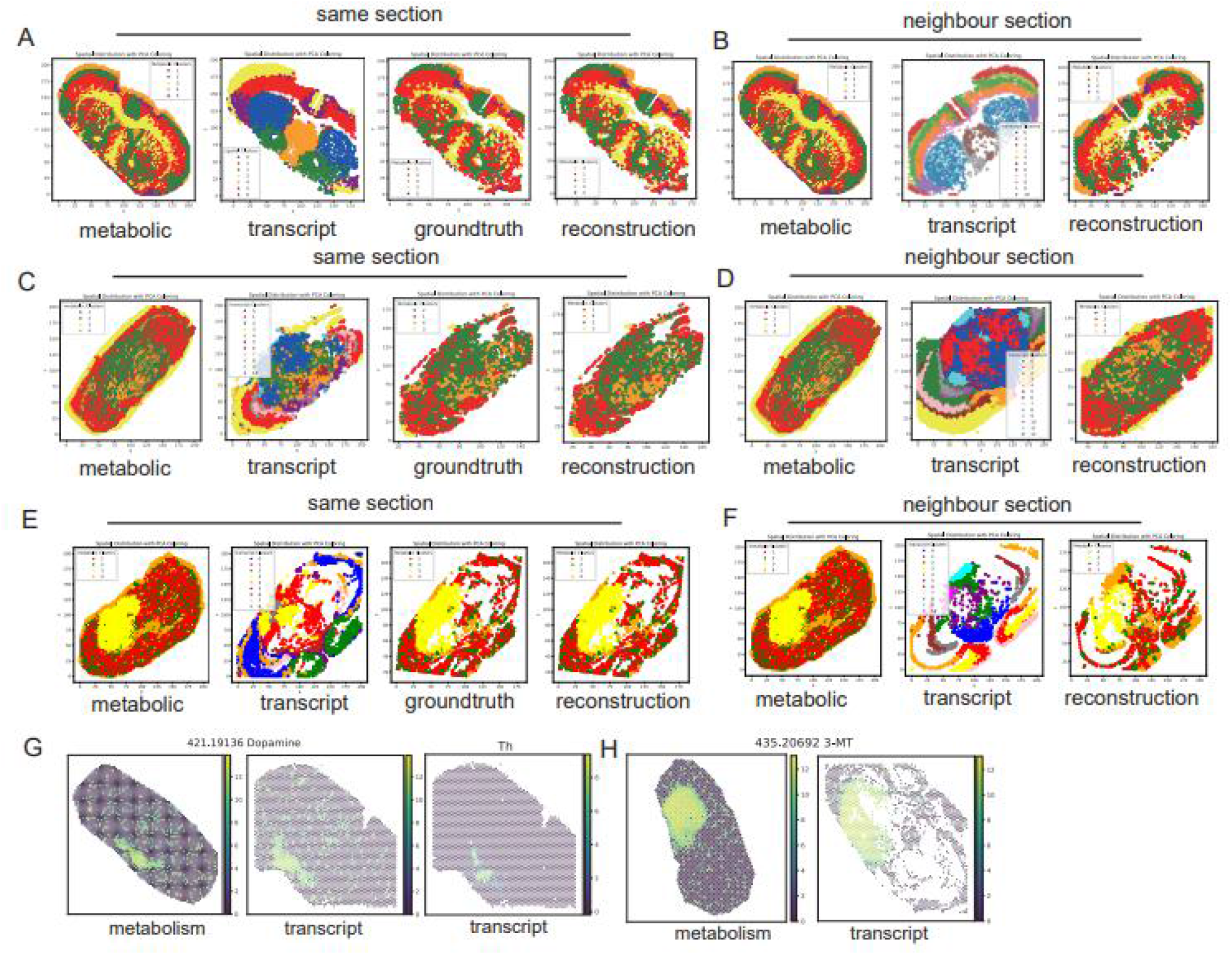
haCCA performs consistent integration of ÎOX Visium and MALDI-MSI spatial data: A-F: spatial plot of MALDI-MSI(metabolic), 10X Visium(transcript), groundtruth and metabolic features of integrated data(reconstruction) of a same coronal section of mouse brain(A,C,E), or neighbour section(B,D,F). G : Dopamine and Dopamine neuron marker Th distribution in mouse MALDI-MSI (sectionl G leftl), spatial transcriptome(section2 G right) haCCA integrating MALDI-MSI to spatial transcriptome(section2 G middle) H: 3-MT distribution in mouse MALDI-MSI (sectionl H left) and haCCA integrating MALDI-MSI to spatial transcriptome(section2 H right)

**Figure 4.**
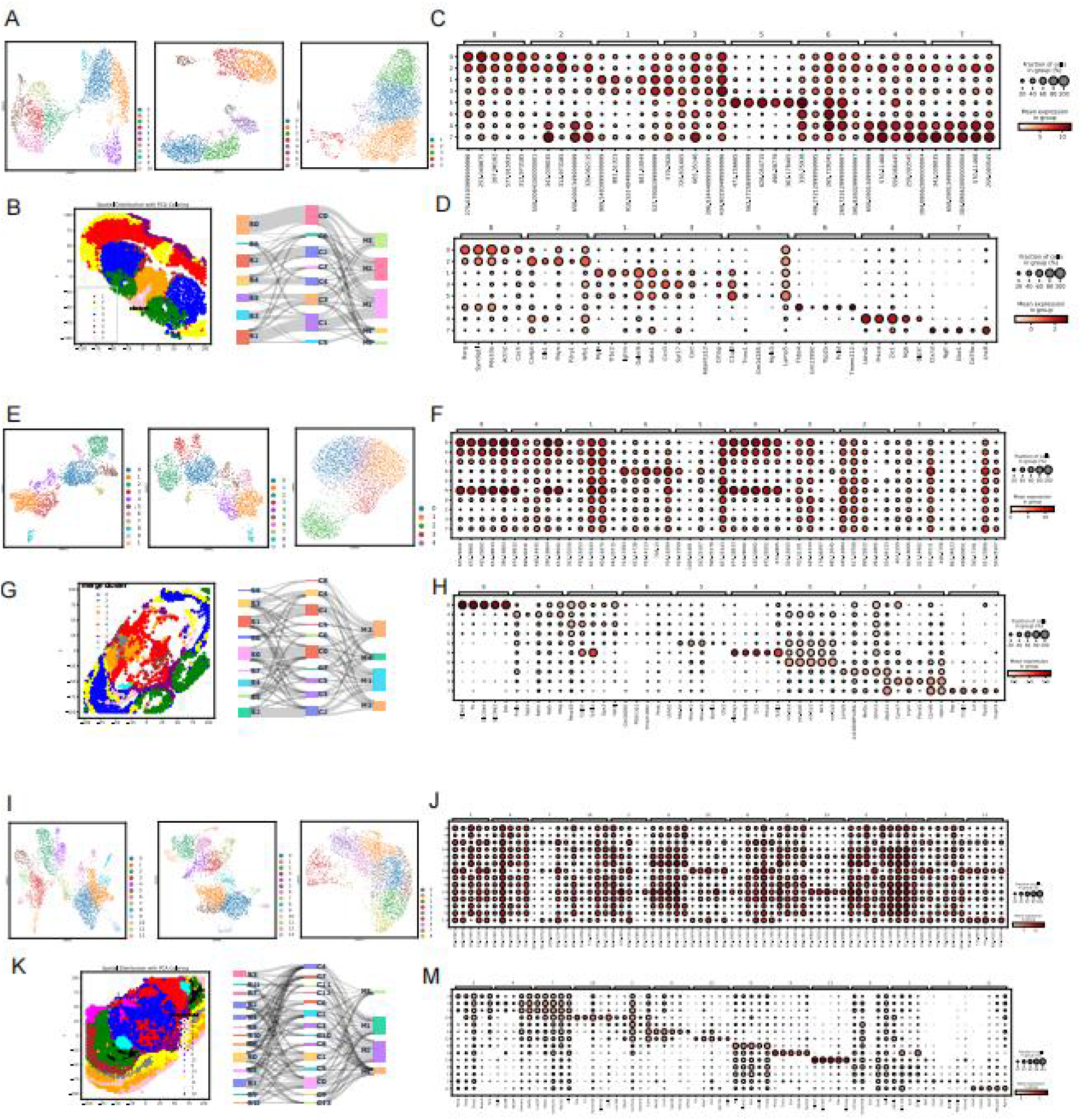
haCCA generated muti-module data allows integrative analysis of transcriptome and metabolome: Umap plot showing cluster of mouse brain using 10X Visium(left), MALDI-MSI(right) and haCCA generated muti-module data(middle)(A,E,l). cluster of haCCA generated muti-module data was projected to 10X Visium spatial plot(B,G,K left). A sanky plot showed the relations of 3 clusters(B,G,K right), marker gene (D,H,M)or metabolites(C,F,J) of clusters in haCCA generated muti-module data of mouse brain.

These results indicated haCCA is able to perform integration between 10X Visium and MALDI-MSI spatial data effectively.

### haCCA generated muti-module data allows integrative analysis of transcriptome and metabolome

After performing haCCA workflow, We performed WNN(weighted nearest neighbors) on the muti-module data contain morphological, transcripts and metabolite data. Compared with using transcript or metabolite data alone, integrated data enable a more dedicate exploration to the heterogeneity of mouse brain by finding more clusters. Different umap plots were shown when using transcript, metabolite or muti-module data for embedding, and a sanky plot shows the relationship between them. Spatially projection was shown to visualize the distribution of integrated data derived clusters(Fig 4A,B,E,G,I,K). We further found the marker gene or metabolites for each cluster(Fig 4 C,D,F,H,J,M). generally, integrated data has the highest heterogeneity, while metabolite data has the lowest. Transcript data derived cluster is highly agreed with the integrated data. These results indicated that haCCA allows integrative analysis of transcript and metabolome data. The heterogeneity of spatial data is mainly contributed by transcripts.

### haCCA generated profile of NETs induced metabolic alteration in preclinical ICC model

Intrahepatic cholangiocarcinoma(ICC) is a type of highly malignant liver cancer. Neutrophil and neutrophil extracellular traps(NETs) consists major components of ICC microenvironment[13, 14]. NETs are a weblike structure extruded by neutrophil, and a few studies revealed its capability to alter the metabolic reprogramming of cancer cells[15]. We eliminated NETs formation in ICC model by knocking out Padi4, and comparing the metabolic alteration of tumor zone by haCCA, to profile the NETs affect on metabolic alteration of ICC. HaCCA showed effective merge of spatial transcriptiome and MALDI-MSI, showing the metabolite on the transcriptome background.(Fig 5A) By eliminating NETs, tumor area of ICC was decreased(1039 spots in KO and 3090 spots in WT, Fig S1), accompanied with previous research(Fig 5B)[16]. We compared the different expressed genes or metabolite (DEG,DEM, scores >15 or < -15 calculated by pyscenic table S2) between WT and KO tumor area, and found in KO groupfwithout NETs), around 73 genes and 77 metabolites were up-regulated, while in WT group(with NETs), around 130 genes and 20 metabolites were up-regulated. KEGG annotation revealed in WT groups, many metabolism associated pathways, such as fatty acid metabolism, cholesterol metabolism and glycolysis or gluconeogenesis was enriched.(Fig 5C) Scd1 is a key enzyme to product monounsaturated fatty acid, and is found to induce tumorigenesis in liver cancer model[17]. Scd1 is the enzyme responsible for oleic acid[18], stearate acid or 18:0 saturated fat acid metabolism[19]. Spatial plot of Scd1, oleic acid, stearic acid and PA(18:0) expression showed a co-localization distribution patternfFig 5D). In cancer area of WT group, those gene/metabolite were also up-regulated compared with KO group.(Fig 5E) We further joint embedded the genes and metabolites in WT group, and found “30 co-expression modules(Table S3). Amongst them we found a cluster consists of ∼45 metabolite and ∼130 genes, centralized by Scd1(Fig S2). Collectively, Using haCCA we perform in vivo and in situ analysis, indicating NETs may alter ICC tumor metabolism by upregulating Scd1. Providing new insights into the dynamic crosstalk between genes and metabolites that regulates the tumor biological behavior.

**Figure 5.**
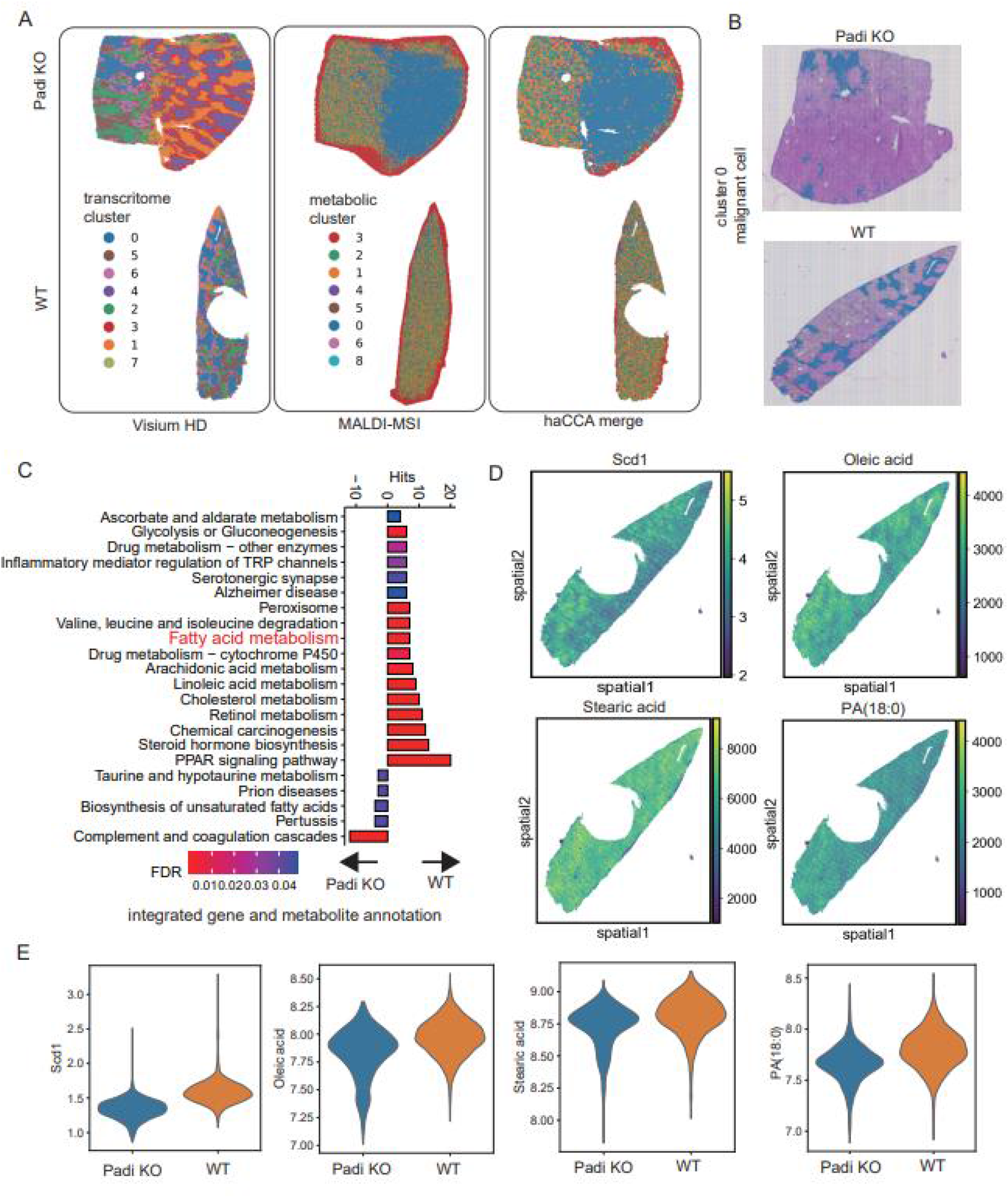
haCCA generated profile of NETs induced metabolic alteration in preclinical ICC model. A : spatial transcriptome(left), MALDI-MSI(middle) and haCCA integrated data(right) of Akt/YapS127A induced ICC model in WT(lower panel) or Padi4 KO(upper panel) mouse. B: malignant cell distribution(Cluster 0) in ICC model. C: KEGG annotation of up-regulation gene and metabolites in malignant cell from WT or KO mouse. D: Scd1, Oleic acid, Stearic acid and PA distribution in ICC from WT mouse. E: Violin plot showed the abundance difference of Scd1, Oleic acid, Stearic acid and PA in malignant cell from KO and WT mouse Figure 6: abstract of haCCA.

## Discussion

Spatial muti-module technologies require efficient and accurate integration of spatial data. Although recently some integration workflow has been developed for spatial transcription data[20-22], their application to muti-module data is limited due to lack of mutual features. Here we proposed haCCA, a method to perform high correlated feature pairs assisted morphological alignment and integration between spatial transcription data and MSI-MALDI data. haCCA is composed of 2 main steps: a modified morphological alignment of spatial coordinates, following by identifying high correlated feature pairs and turning them into new high correlated variables. Using aligned spatial coordinates and high correlated variable to perform integration with more accuracy than only using morphological alignment.

By using spatial transcription data and pseudo paired MSI-MALDI data, we shown that haCCA outperformed morphological alignment by 6-20%, represented by STUtility, evaluating by label transfer accuracy, label transfer ARI. Besides, the modified morphological alignment of spatial coordinates of haCCA itself is able to outperform STUtility, and the combing alignment is able to further increase the accuracy. It indicated the efficiency of combing alignment strategy of haCCA. When using real world generated data of normal tissue and tumor, haCCA successfully integrated MSI-MALDI data into spatial transcriptome data, and generated gene expression as well as bio molecule profiling. Such multi-module spatial data allows more sophisticated deciphering to heterogeneity, for example, we identified different metabolic states in tumor region from ICC model. It also facilitates both gene and bio molecule marker detection for certain clusters.

haCCA may be improved in the following aspects. First, only spatial transcription and spatial metabolome data has been used. haCCA can expend its capability to integrate more spatial techniques, such as image-based sequencing, spatial proteinomes or histological images. Second, how haCCA behaves on higher resolution data should be further explored.

## Conclusion

To summarize, we presented a workflow named haCCA that enables the simultaneous spatial profiling of metabolites and transcriptome across neighbor tissue section. Applying haCCA to tumor samples could increase understanding of both metabolic and transcript heterogeneous, providing new insights into the dynamic crosstalk between genes and metabolites that regulates the tumor biological behavior and drives the response to treatment.(Figure 6)

**Figure 6.**
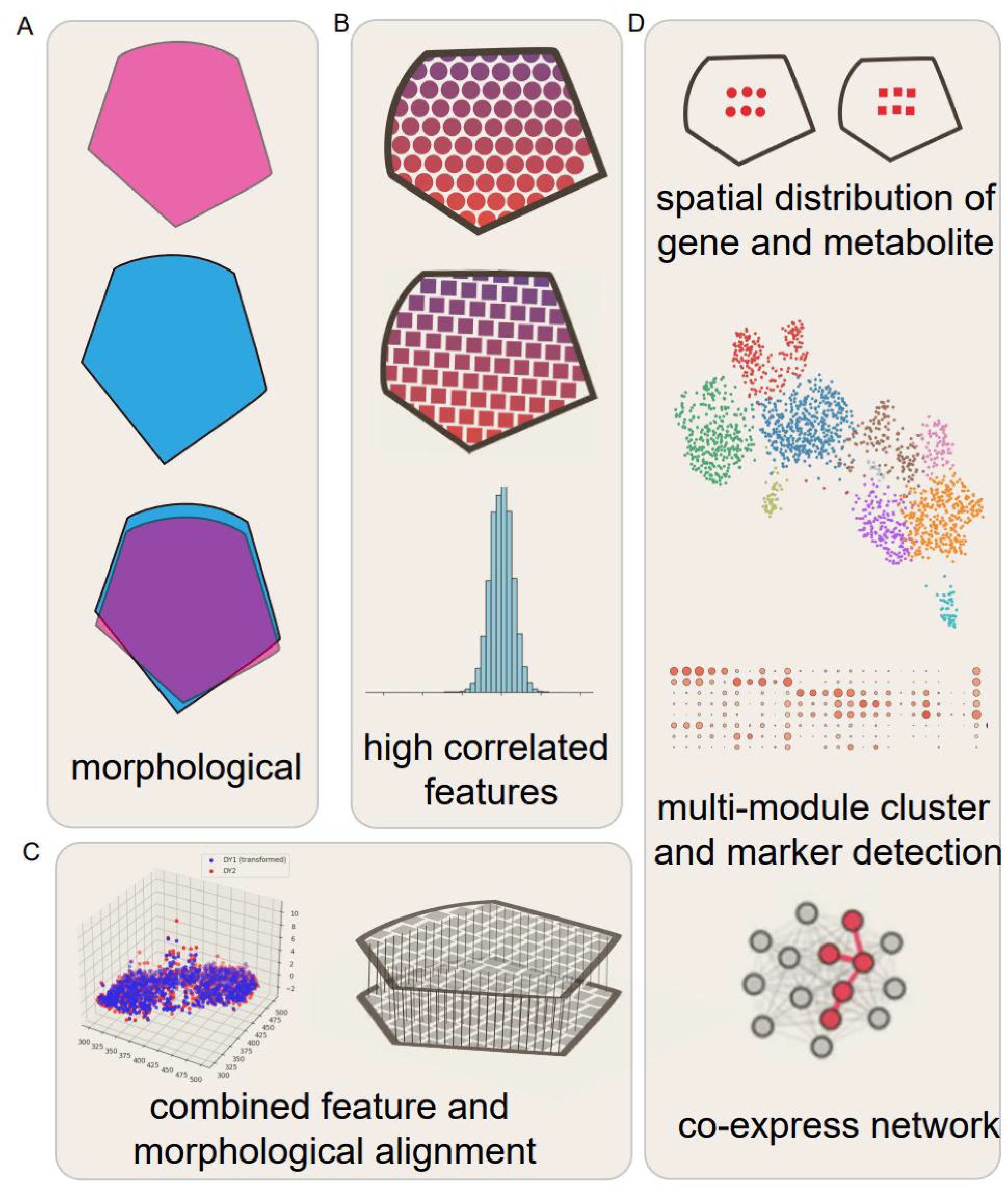
abstract of haCCA.

## Figure legend

**Figure S1:**
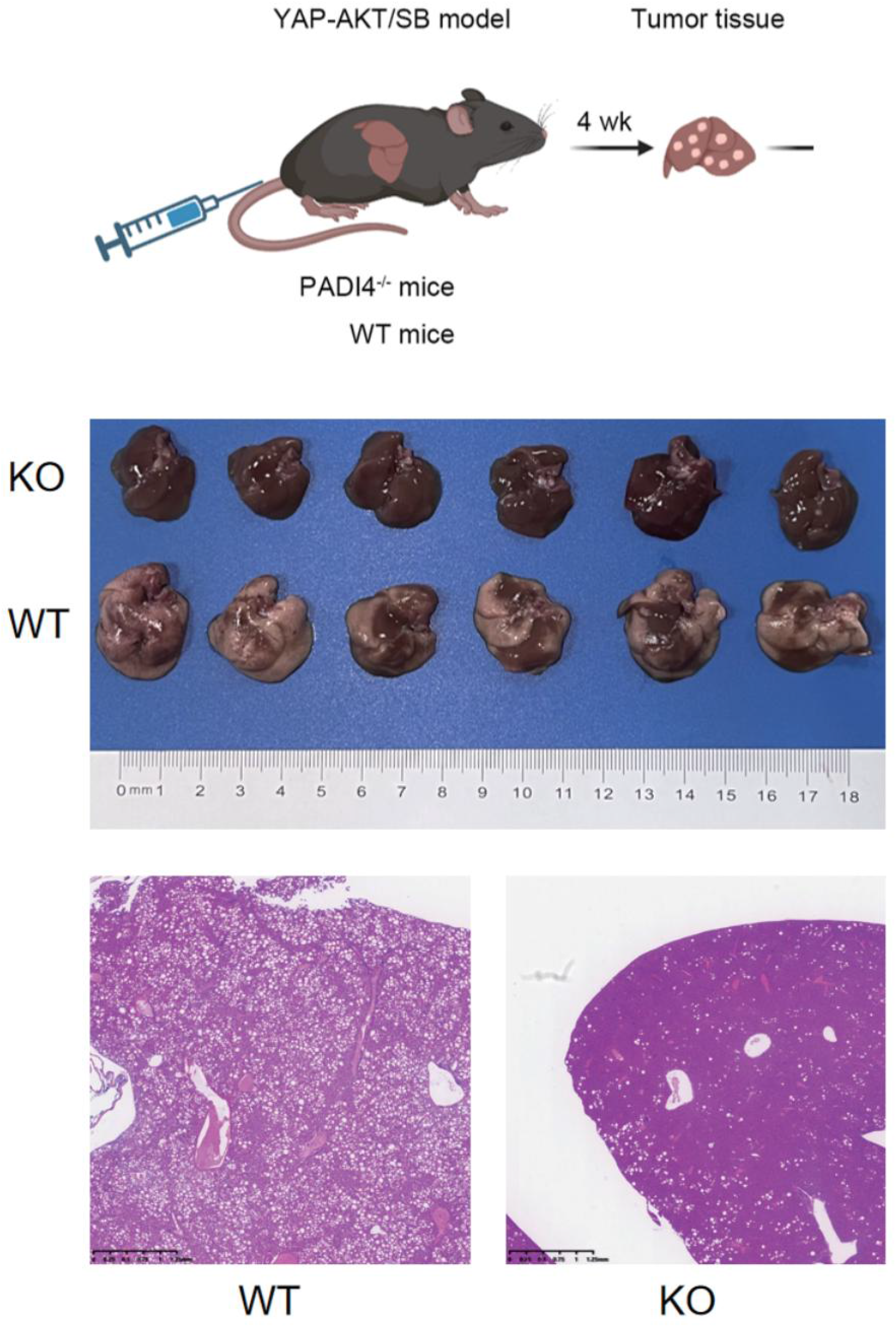
establishment of ICC mouse model. Upper: scheme plot of model establishment. Padi4 KO or WT C57 mouse were HDI injected with lOug plasmid. 4w later tumor were collected for further experiment. Middle: gross appearance of KO or WT tumor. Below: HE staining of KO or WT ICC tumor

**Figure S2:**
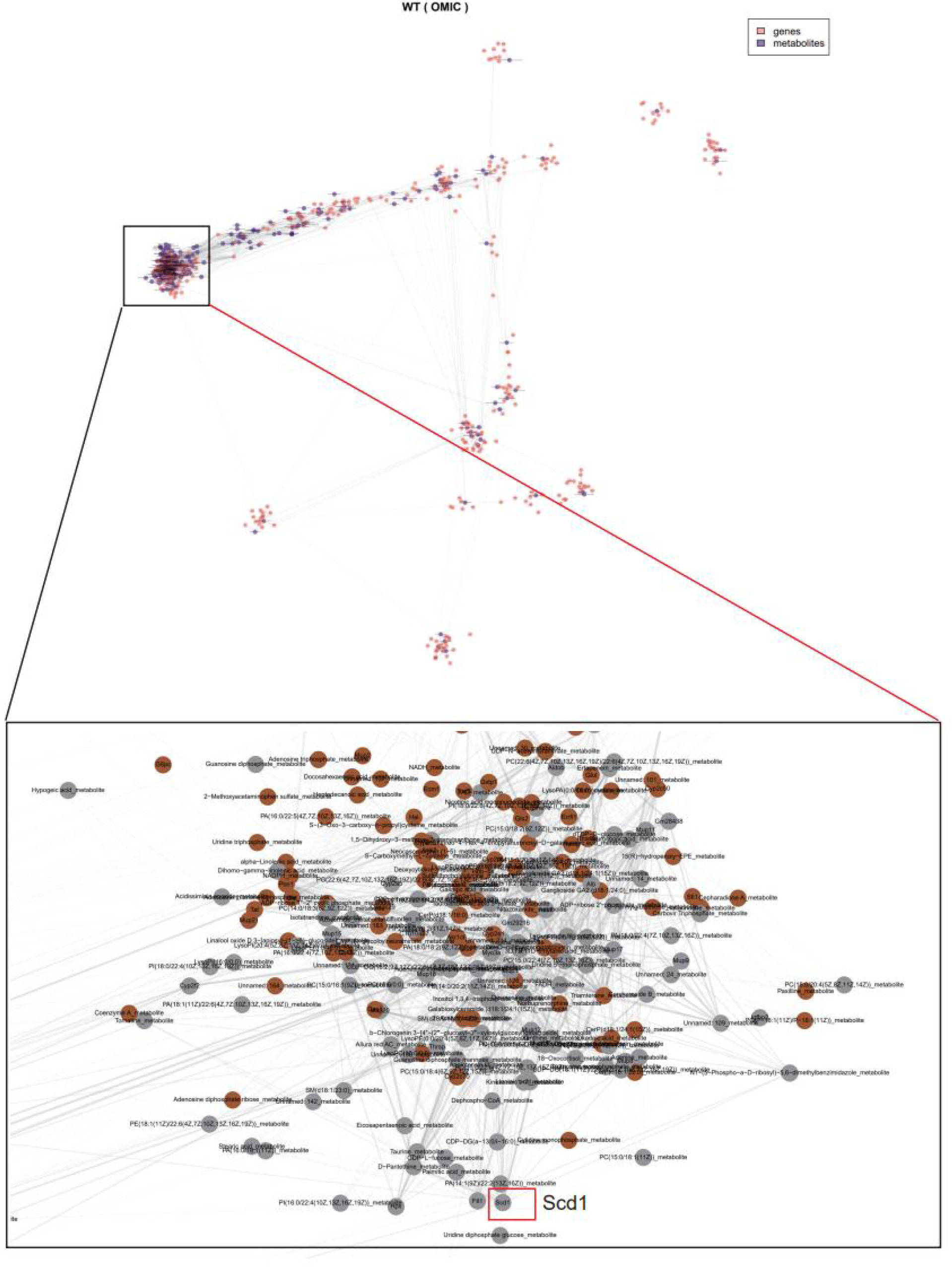
joint embedding of genes and metabolites in WT ICC. Nodes (genes and metabolites) were split in 30 spatially co-detected modules using the Spinglass algorithm.

## Declarations

### Ethics approval and consent to participate

The present study was performed in accordance with the Declaration of Helsinki. Approval for the use of human subjects was obtained from the research ethics committee of Huashan Hospital, Fudan University(KY2023-594), and informed consent was obtained from each individual enrolled in this study. All animal experiments were approved by the Animal Ethics Committee of Fudan University

### Consent for publication

Not applicable

### Availability of data and materials

The datasets used and/or analysed during the current study are available from the corresponding author on reasonable request.

### Code availability

The code and example of haCCA is available at GitHub LittleLittleCloud/haCCA: anchor spots pairs and high correlated feature pairs assist multi-module spatial assay integrating

## Acknowledgements

We sincerely thank Dr Chen-De Yang and Dr Dan-Ye in assisting establishing mouse model, Dr. Yang Zhang in assisting coding and writing improvement.

